# Temporal epigenome modulation enables efficient bacteriophage engineering and functional analysis of phage DNA modifications

**DOI:** 10.1101/2024.01.28.577628

**Authors:** Nadiia Pozhydaieva, Franziska Anna Billau, Maik Wolfram-Schauerte, Adán Andrés Ramírez Rojas, Nicole Paczia, Daniel Schindler, Katharina Höfer

**Author notes:** Corresponding author: Katharina Höfer, PhD.

## Abstract

Lytic bacteriophages hold substantial promise in medical and biotechnological applications. CRISPR-Cas systems offer a way to explore these mechanisms via site-specific phage mutagenesis. However, phages can resist Cas-mediated cleavage through extensive DNA modifications like cytosine glycosylation, hindering mutagenesis efficiency. Our study utilizes the eukaryotic enzyme NgTET to temporarily reduce phage DNA modifications, facilitating Cas nuclease cleavage and enhancing mutagenesis efficiency. This approach enables precise DNA targeting and seamless point mutation integration, exemplified by deactivating specific ADP-ribosyltransferases crucial for phage infection. Furthermore, by temporally removing DNA modifications, we elucidated the effects of these modifications on T4 phage infections without necessitating gene deletions.

Our results present a strategy enabling the investigation of phage epigenome functions and streamlining the engineering of phages with cytosine DNA modifications. The described temporal modulation of the phage epigenome is valuable for synthetic biology and fundamental research to comprehend phage infection mechanisms through the generation of mutants.

## Main

Bacteriophages (phages) are viruses that specifically infect prokaryotic hosts. The high potential of the application of lytic phages in both medical and industrial settings has boosted phage research in recent years^1–4^. However, a comprehensive understanding of the molecular mechanisms that underlie efficient phage infections, which is essential for their targeted utilization, remains significantly underexplored. This knowledge gap is evident even in extensively studied phages like the *Escherichia coli* bacteriophage T4 (phage T4). Approximately half of the 273 encoded proteins in phage T4 are associated with known functions, while the roles of the other proteins remain unclear^5^. To explore the biological functions of phage-derived proteins, gene deletion has been the primary method historically employed, providing critical insights into gene essentiality for the phage infection process^6,7^. However, mutagenesis targeting only the catalytic residues rather than deleting the entire protein offers a possibility to elucidate the enzyme’s role within a specific molecular context. The generation of catalytically inactive proteins preserves the other potential functions of the studied protein, such as involvement in protein-protein or protein-nucleic acid interactions. Particularly in phages with complex genomes, like phage T4, which has numerous overlapping genes and gene splicing arrangements^7^, targeted mutagenesis is crucial to avoid unintended effects on other genes’ expression.

Prophages, whose genomes are integrated into the host chromosome and passively replicate alongside the host genome, can be mutated *in vivo* using the same genetic tools as for the mutagenesis of their host. However, this does not apply to lytic phages, as their genetic material exists separately from the host genome within bacterial cells and undergoes rapid replication during a limited time of infection. Therefore, targeted mutagenesis of lytic phages is highly challenging, as evidenced by the diverse strategies developed in recent years to approach it (reviewed in Mahler *et al.* 2023 ^8^). Many developed approaches rely on recombination between phage DNA and a donor sequence, e.g., homologous recombination. This approach allows for gene replacements, deletions, or insertions. However, the efficiency of this method is low (>0.05%), resulting in the requirement of extensive screening to identify phage mutants^9,10^. To streamline the screening, incorporating reporter genes alongside the mutation to simplify the detection of the mutant phages has been employed^10,11^. Yet, the insertion of reporter genes may affect the complex gene expression of the phage, like the implications of phage gene deletions as described above. Additionally, inserting an extra gene may affect the packaging of phage DNA into the capsid, which has a fixed size^10^.

Another approach, building on homologous recombination principles, is the bacteriophage recombineering of electroporated DNA (BRED) method, which offers a relatively high mutagenesis rate (10-15%)^12^. However, its effectiveness depends on the successful co-transformation of both phage DNA and donor DNA into the same cell, which might be limited by the host’s transformation efficiency or the phage genome size^13^.

The discovery of the antiphage clustered regularly interspaced short palindromic repeats (CRISPR) and the CRISPR-associated protein (Cas) has revolutionized the field of genome editing. The successful application of targeted CRISPR-Cas-based mutagenesis across various organisms^14^, has also raised the interest in its application for phage engineering. In this context, CRISPR-Cas can be employed to target a specific position within the phage genome during infection, generating a DNA double-strand break. This break can be further repaired via homologous recombination with donor DNA (DNA carrying the desired mutation) present within the infected cell. However, the application of the CRISPR-Cas-based phage mutagenesis on the model phages, such as phage T4, has revealed an overall low mutagenesis efficiency (0-3%) and being strongly dependent on the spacer selected for the mutagenesis^15–17^. This strongly impedes the specific targeting of the phage genome via CRISPR-Cas and requires pre-screening for efficient spacer before the mutagenesis, strongly reducing the applicability of the approach for efficient phage mutagenesis.

The reason for the strong spacer dependence of Cas targeting efficiency is attributed to the extensive DNA modifications - present on phage DNA - that protect it from cleavage. The DNA modifications among the phages are widely distributed and are also present in the phage T4^18,19^. The enzyme deoxycytidylate 5-hydroxymethyltransferase (*gene 42*) originating from T4 phage catalyzes the conversion of 2′-deoxycytidine 5′-monophosphate (dCMP) into 5-hydroxymethyl-2′-deoxycytidine 5′-monophosphate, which is subsequently converted to into 5-hydroxymethyl-2′-deoxycytidine 5’-triphosphate and incorporated into T4 DNA during replication (further referred to as 5hmdC). Next, the 5hmdC is glycosylated by α- or β-glycosyltransferases (α-/β-gt) to 5-α-/β-glycosyl hydroxymethyl-2′-deoxycytidine (5ghmdC) within T4 DNA (Fig. 1a)^20,21^. The T4 phage epigenome plays a crucial role in phage fitness, exemplified by the essential *gene 42*. Notably, amber mutants for the gene have been reported. However, in this scenario, phage infections must be carried out using *E. coli* strains that harbor additional plasmids and thus do not represent the wild-type *E. coli*. Furthermore, *α-/β-gt* serve as auxiliary genes^7,17,22^, as the encoded proteins are pivotal in protecting phage DNA through glycosylation from host defense systems and host nucleases, including Cas nucleases^15,23^. Thus, DNA modifications such as 5hmdC and 5ghmdC prevent the effective use of CRISPR-Cas for targeted phage mutagenesis and counterselection.

**Fig. 1:**
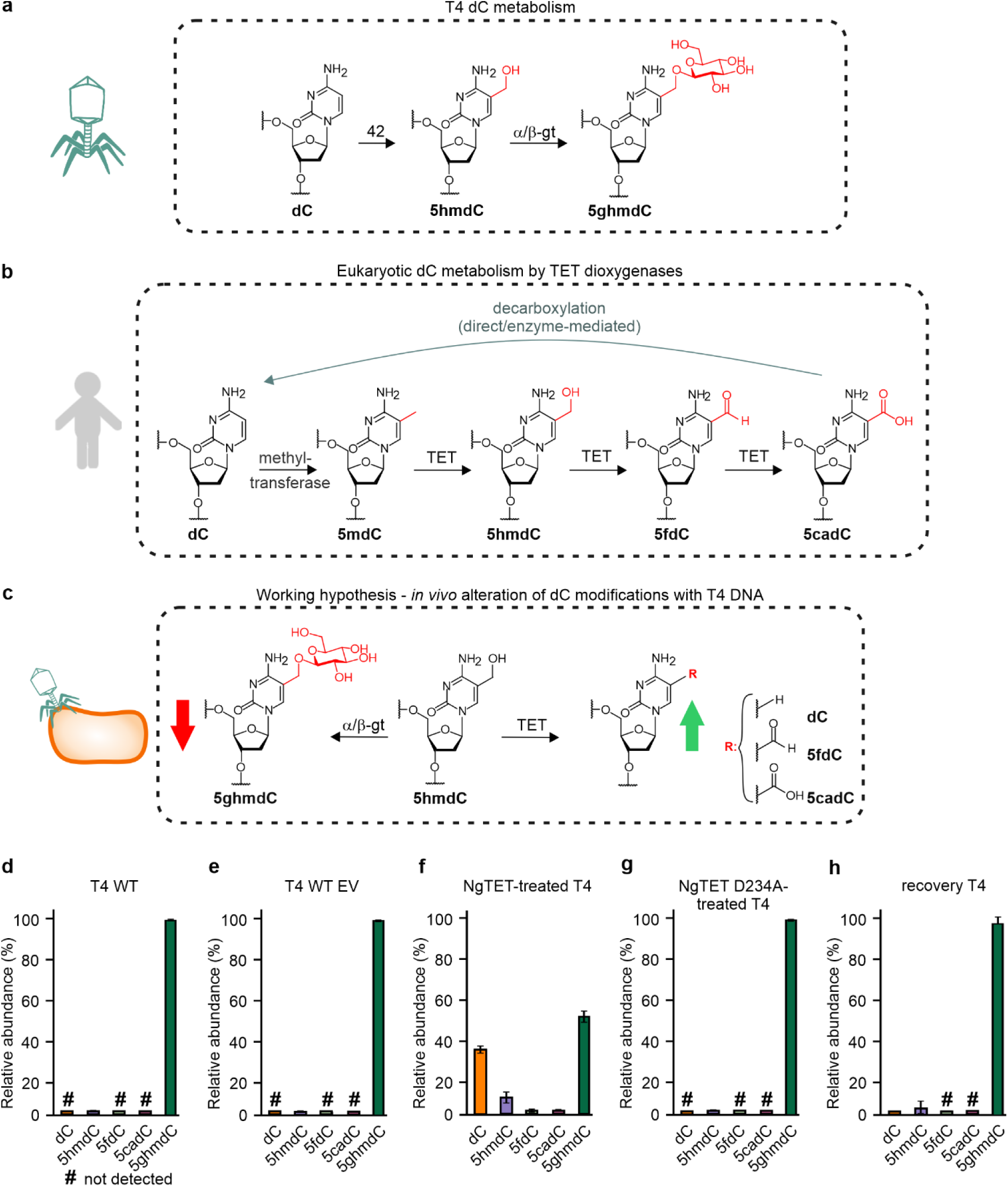
T4 phage DNA is extensively modified. **a**, phage T4 DNA is extensively modified by deoxycytidylate 5-hydroxymethyltransferase (42) (oxidation of dC to 5hmdC), and α/β-gt (glycosylation of 5hmdC to 5ghmdC). **b,** TET dioxygenase plays a crucial role in eukaryotic epigenetic regulation by demethylating 5mdC through its iterative oxidation. **c,** 5hmdC present in phageT4 is one of the natural substrates of TET. Therefore, the glycosylation of 5hmdC to 5ghmdC by α-gt/β-gt is expected to be downregulated in the presence of TET due to substrate competition between TET and α-gt/β-gt. **d-h,** Relative abundance (%) of 2′-deoxycytidine metabolites determined via LC-MS analysis in different T4 strains: T4 WT (**d**), T4 WT propagated in the presence of empty vector (EV) (T4 WT EV) (**e**), NgTET-treated T4 (**f**), NgTET D234A-treated T4 (**g**), and recovery T4 (**h**). The presence of dC traces in recovery T4 DNA (**h**) may be attributed to residual NgTET-treated T4 phages that did not infect *E. coli* and therefore were not recovered. Hashtag highlights the nucleosides not detected in a sample. *n =* 3 biological replicates. The significance of specific cytosine modification changes is shown in Extended Data Fig. 2e.

In this study, we apply a eukaryotic ten-eleven translocation (TET) methylcytosine dioxygenase^24^ to temporally reduce the abundance of 5ghmdC within the phage T4 genome to enable the specific and efficient targeting of phage DNA with CRISPR-Cas. This results in the facilitated introduction of point mutations into the phage genome in a scarless manner, exemplarily shown for two specific T4 phage ADP-ribosyltransferases crucial for phage infection. The increased targeting efficiency of phage DNA by Cas nucleases allows for sequence-specific spacer selection. Simultaneously, the improved targeting efficiency of phage DNA by Cas nucleases can be exploited for efficient counterselection, streamlining the enrichment process for the generated phage mutant. We utilized a high-throughput screening approach based on next-generation sequencing (NGS) to identify the mutants, facilitating the identification and validation of phage T4 mutants. This method is compatible with automation, simplifying the phage genome engineering process and eliminating the need to introduce reporter genes into the phage genome. The scarless nature of this approach allows for the precise study of the impact of introduced mutations on phage infection without the side effects associated with gene deletion or reporter gene insertion. Overall, our findings propose an efficient strategy for introducing point mutations into genomes of lytic phages that possess cytosine-based DNA modifications. These advantages collectively position this technique as a valuable tool in synthetic biology and biotechnology for creating "designer phages" as well as in fundamental research to explore the mechanisms underlying phage infections. Moreover, this technique offers the opportunity to study the influence of phage epigenome on phage infection dynamics without necessitating the deletion of the essential phage DNA modifying genes.

## Results

### Eukaryotic NgTET modulates the abundance of T4 DNA modifications

Phage DNA modifications, such as 5ghmdC, impede the recognition and targeting of phage DNA by host nucleases and CRISPR-Cas systems^25^. The absence of these modifications would enhance phage DNA accessibility to host nucleases, resulting in phage genome degradation and the prevention of successful propagation^26,27^. No host-or phage-originating enzymes are currently known to specifically remove 5ghmdC modifications from the phage T4 genome.

In contrast to the 5ghmdC modification, enzymes that act on its direct precursor, 5hmdC, are well-known within the eukaryotic realm and play crucial roles in epigenetic regulations. Eukaryotic TET dioxygenases are involved in active DNA demethylation via iterative oxidation of 5-methyl-2′-deoxycytydine (5mdC) to 5hmdC, 5-formyl-2′-deoxycytidine (5fdC), and 5-carboxy-2′-deoxycytidine (5cadC) (Fig. 1b)^28,29^. In the next step, 5cadC undergoes non-enzymatic decarboxylation, leading to the formation of unmodified 2′-deoxycytidine (dC)^30,31^. Therefore, we hypothesized that eukaryotic TET enzymes could potentially be harnessed to convert 5hmdC modifications in phage DNA into 5fdC and 5cadC (Fig. 1c). Leveraging the demethylation activity of TET enzymes on phage DNA presents an opportunity to diminish 5hmdC levels and compete effectively with T4 phage α/β-gt enzymes for the substrate 5hmdC during T4 phage infection. This approach has the potential to decrease the prevalence of 5ghmdC in phage DNA, thereby enhancing the efficacy of CRISPR-Cas-based editing, as depicted in Figure 1c.

To test this hypothesis, we selected the well-characterized TET enzyme from the single-cell protist *Naegleria gruberi* (NgTET), which has already been expressed in *E. coli* in a catalytically active form^32,33^. First, we evaluated the influence of heterologous NgTET expression on *E. coli* growth to confirm that the dioxygenase expression does not impede host growth and, consequently, does not affect the host’s susceptibility to phage infection (see Extended Data Fig. 1a-b). Notably, the expression of NgTET in *E. coli* did not significantly impact the bacterial growth rate or bacterial lysis by T4 phage. Moreover, phages derived from the lysis of the NgTET-expressing strain were still found to be infectious and cause complete lysis when infecting wild-type *E. coli* (Extended Data Fig. 1b).

Next, we investigated if NgTET is active on phage T4 DNA. For this, we evaluated if the expression of NgTET in *E. coli* during phage T4 infection alters the presence of 2′-deoxycytidine modifications such as 5hmdC and 5ghmdC in the phage DNA. To do this, we infected an *E. coli* strain expressing NgTET with T4 wild-type (T4 WT) phages. We conducted a comprehensive LC-MS-based analysis of the DNA composition of the resulting T4 phages, which we refer to as "NgTET-treated T4" phages. This analysis enabled us to compare the DNA composition of the phage progeny obtained from the NgTET-expressing *E. coli* strain, NgTET-treated T4, with that of the phage progeny resulting from the infection of *E. coli* wild-type cells, T4 WT (see Figs. 1d-g and Extended Data Fig. 2). In agreement with previous studies that characterize the T4 WT DNA composition^21^, we observed the complete absence of unmodified 2′-deoxycytidine in T4 WT DNA. According to our LC-MS analysis, >99% (expressed as relative abundance (r.a.)) of all 2′-deoxycytidines were present as 5ghmdC and <1% - as 5hmdC (Fig. 1d-e).

The analysis of NgTET-treated T4 DNA revealed a reduction of 5 ghmdC from 99% r.a. (T4 WT) to 55% r.a. (NgTET-treated T4) (Fig. 1f). In addition, 5hmdC (10.5% r.a.), 5fdC (2.3% r.a.), 5cadC (0.9% r.a.), and unmodified 2′-deoxycytidines, dC (34.4% r.a.) were detected in the NgTET-treated T4 DNA (Fig. 1b). 5fdC and 5cadC are known oxidation products of 5hmdC^32,33^ (Fig. 1b). The significant reduction in 5ghmdC levels (Extended Data Fig. 2e), coupled with the presence of unmodified dC, which is the result of the decarboxylation of 5cadC^30,31^, confirms the catalytic activity of NgTET on T4 DNA.

To unambiguously link the altered modifications abundance in T4 DNA with NgTET enzymatic activity, we generated a catalytically inactive NgTET D234A mutant as a negative control^34^. LC-MS analysis of DNA isolated from the phage progeny released from *E. coli* strain overexpressing inactive NgTET D234A (referred to as "NgTET D234A-treated T4") showed a DNA composition similar to that of T4 WT (Fig. 1g) (>1% r.a. for 5hmdC and 99% r.a. for 5ghmdC in NgTET D234A-treated T4 DNA). These findings make us confident that the observed reduction of 5ghmdC is due to the catalytic activity of NgTET, expressed in *E. coli* during T4 phage infection, efficiently oxidizing 5hmdC within phage T4 DNA.

### Modulation of phage DNA modifications by NgTET alters bacterial cell lysis

As emphasized earlier, DNA modifications significantly contribute to phage fitness, and their absence can potentially alter the phage infection phenotype. Based on this, we aimed to investigate whether the phenotype of NgTET-treated T4 phages might be altered due to reduced abundance of 5ghmdC modification. Specifically, we hypothesized the host cell lysis being affected for NgTET-treated T4 compared to the lysis by T4 WT phages, as less modified phage DNA would be more susceptible to degradation by host nucleases (Fig. 2a). Consequently, the replication and gene expression machinery of the phage will be impeded, affecting the lysis of the bacterial host.

**Fig. 2:**
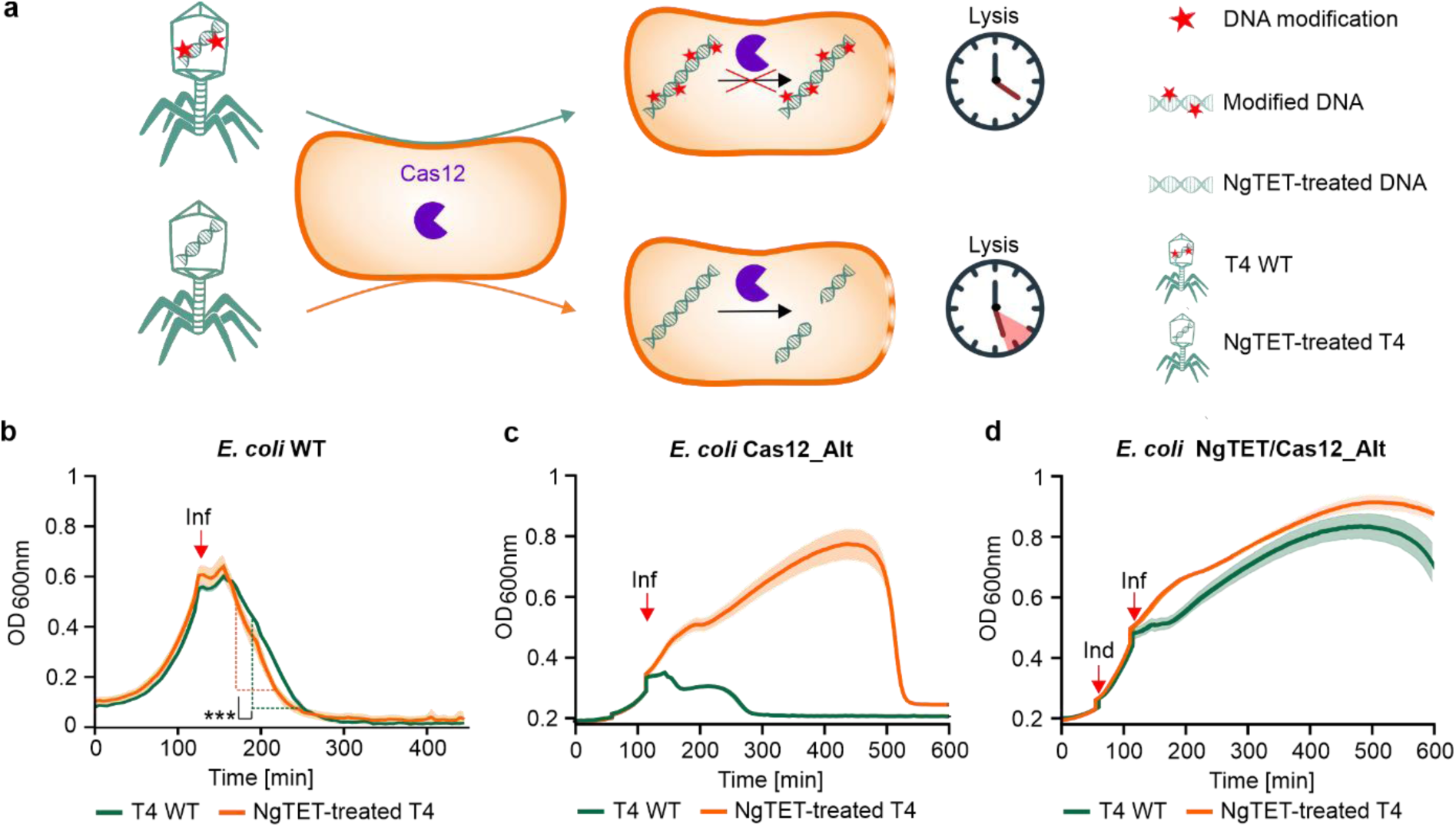
Impact of T4 DNA modifications on T4 phage lysis behaviour in the presence and absence of CRISPR-Cas12. **a**, T4 DNA modifications protect phage DNA against Cas12 nuclease targeting, ensuring efficient phage propagation cell lysis. Reduced DNA modifications increase Cas12 nuclease susceptibility, leading to impeded and therewith delayed lysis of bacterial culture. **b**, Lysis kinetics of *E. coli* infected by T4 WT and NgTET-treated T4. Red arrow highlights the time point of the phage addition (Inf) (two-sided Student’s t-test, *P* = 0.0096 at *P*_signif_ <0.05). *n =* 3 biological replicates. **c**, Lysis kinetics of *E. coli* Cas12_Alt by T4 WT and NgTET-treated T4. Red arrows highlight the time point of NgTET expression induction (Ind) and the addition of the phage (Inf). *n =* 3 biological replicates. **d**, Lysis kinetics of *E. coli* NgTET/Cas12_Alt by T4 WT and NgTET-treated T4. Red arrows highlight the time point of NgTET expression induction (Ind) and the addition of the phage (Inf). *n =* 3 biological replicates.

To answer this question, we infected wild-type *E. coli* with either T4 WT or NgTET-treated T4 (Fig. 2b) and monitored bacterial cell lysis. In line with our hypothesis, NgTET-treated T4 phages exhibited a notably slower lysis rate than the T4 WT phages, resulting in an approximately 15-minute delay in the onset of lysis. Nevertheless, both T4 WT and NgTET-treated T4 phages ultimately induced complete lysis of the bacterial culture. Furthermore, the phages released during the lysis with NgTET-treated T4 were confirmed to be infectious, as evidenced by their ability to infect and lyse wild-type *E. coli* (Extended Data Fig. 1b).

### Temporal removal of DNA modifications enhances CRISPR-Cas12 targeting of phage DNA

The previous experiment showed that treating phage T4 with NgTET led to diminished lysis of *E. coli*, showing the importance of phage DNA modifications for its fitness. Building on this, we aimed to investigate whether CRISPR-Cas targeting of the phage DNA would also be enhanced due to the NgTET treatment. We hypothesized that the reduction in DNA modifications could increase the susceptibility of phage DNA to Cas nuclease targeting, as demonstrated by the more effective targeting of non-glycosylated phage T4 DNA by CRISP-Cas12^35^. Therefore, NgTET treatment of T4 DNA would potentially result in Cas-mediated DNA double-strand breaks. Such a break would impede phage replication, consequently causing delayed or even absent lysis by NgTET-treated T4 compared to T4 WT. Given the extensive modifications in T4 WT DNA, we expected its lysis not to be significantly delayed by the expression of the CRISPR-Cas system.

To follow this hypothesis, we conducted a lysis experiment in which we infected an *E. coli* strain, heterologously expressing the CRISPR-Cas12, with either T4 WT or NgTET-treated T4 phages (Fig. 2c). For a proof of principle study, we designed a spacer targeting the T4 gene *alt* which encodes the T4 Alt ADP-ribosyltransferase. First, we verified that overexpression of CRISPR-Cas12 without a spacer does not negatively affect phage lysis and bacterial growth (Extended Data Fig. 3). Next, we conducted the infection of the *E. coli* strain expressing the CRISPR-Cas12_Alt (spacer targeting *alt* gene) with T4 WT. As expected, we did not observe any negative impact of CRISPR-Cas12_Alt expression on phage lysis. This observation demonstrates that DNA modifications protect the genome of phage T4 from nucleolytic cleavage by Cas12, enabling efficient phage lysis.

Next, we analyzed the lysis by NgTET-treated T4 in the same experimental settings. While infecting *E. coli* expressing CRISPR-Cas12_Alt, a delay of approximately 250 min in lysis compared to infection with T4 WT was observed. This delay supports our hypothesis that the reduced abundance of 5ghmdC in NgTET-treated T4 enables Cas nuclease targeting of phage DNA. Consequently, phage propagation is impeded, which is reflected in the delayed onset of lysis.

To confirm that this delay is attributed to the targeting of T4 DNA by CRISPR-Cas12_Alt, we infected *E. coli* expressing the CRISPR-Cas12 without any spacer, thus lacking any target, with either T4 WT or NgTET-treated T4. Since there was no spacer in CRISPR-Cas12, no targeting of phage DNA and no additional impact on phage lysis was expected. Consistent with our expectations, we observed the same lysis behaviour for NgTET-treated T4 infection of the CRISPR-Cas12 expressing *E. coli* strain as for the infection of wild-type *E. coli* (Extended Data Fig. 1d). This confirms that the observed delay in Fig. 2c is a direct consequence of CRISPR-Cas12_Alt efficiently targeting the *alt* gene in DNA of NgTET-treated T4 phage.

### Continuous removal of T4 DNA modifications prevents bacterial lysis

Despite the observed lysis delay in *E. coli* expressing CRISPR-Cas12_Alt when infected by NgTET-treated T4 phages in the previous experiment, the eventual lysis of the culture indicates that the phages ultimately succeeded in propagating efficiently (Fig. 2c). We hypothesized, that NgTET-treated T4 phages regain their lysis capacity during infection. We speculated that the progeny of NgTET-treated T4 phages, which evaded CRISPR-Cas12_Alt targeting, restored wild-type-like DNA modification levels, as NgTET dioxygenase was not heterologously expressed upon infection.

To prove our hypothesis, we conducted a further lysis experiment. This time, we infected *E. coli* cells that were simultaneously expressing both CRISPR-Cas12_Alt and NgTET dioxygenase. The constant overexpression of the NgTET was aimed to maintain a reduction in DNA modifications in the phage progeny generated during the experiment. Notably, upon expression of NgTET dioxygenase, we did not observe the onset of lysis for up to 600 min post-infection in both T4 WT and NgTET-treated T4 infections (Fig. 2d), which strongly supports our hypothesis.

In conclusion, these findings underscore the critical role of T4 DNA modifications in protecting the DNA from host anti-phage defense systems, particularly those operating at DNA level, such as CRISPR-Cas12. While the reduction in 5ghmdC levels in NgTET-treated T4 seems to enhance phage DNA susceptibility to CRISPR-Cas12, it simultaneously leads to decreased phage fitness and a substantial alteration of the phage phenotype (Fig. 2d).

### The removal of DNA modifications by NgTET is reversible

To preserve the integrity of phage phenotypes in genetic studies, particularly when utilizing phage strains with decreased DNA modifications for mutagenesis, it is crucial to restore wild-type (WT) DNA modification levels. In this study, we aimed to temporally alter T4 DNA modification levels during mutagenesis and restore them to T4 WT levels after successfully introducing the mutation of interest. In such a way, only the impact of the introduced mutations can be studied. The reversible alteration of the T4 epigenome represents a notable advantage compared to employing Δα-/β-gt T4 or respective amber T4 mutants, where permanent alterations of phage DNA modifications can affect the phage phenotype^22^.

We hypothesized that treating T4 DNA with NgTET could be used to decrease 5ghmdC levels temporarily. Hence, our investigation aimed to determine if the reduction in the 5ghmdC fraction within the NgTET-treated T4 DNA, is reversible to wild-type levels in the subsequent phage generation, termed as "recovery T4", produced in the absence of NgTET dioxygenase.

To confirm that the effects of phage DNA treatment with NgTET are indeed reversible and do not impact DNA modifications in the subsequent progeny, we infected *E. coli* WT with NgTET-treated T4. The DNA composition of recovery T4 was analyzed by LC-MS (Fig. 1h). Our data show that already for the first generation of recovery T4 phages, the modification levels were comparable to the modification levels in T4 WT DNA (3% r.a. for 5hmdC, 97% r.a. for 5ghmdC, and traces of >0.3% dC). The absence of the NgTET-mediated oxidation products, 5fdC, and 5cadC, confirms that the altered abundance of DNA modifications in NgTET-treated T4 is transient and can be restored in the next phage generation.

Next, the recovery phage was also analyzed regarding its infectivity of the host, demonstrating same lysis rate as T4 WT phage (Extended Data Fig. 1b).

### NgTET treatment of T4 DNA allows efficient Cas-mediated phage DNA targeting *in vivo*

To assess the impact of NgTET treatment of phages on Cas-mediated genome engineering efficiency, we selected two T4 phage-encoded genes, *modA,* and *alt*, as mutation targets. Alt and ModA belong to the enzyme class of ADP-ribosyltransferases. During infection, they have been described to play a crucial role in the takeover of control over *E. coli* cells by T4 phage, although the exact mechanisms behind this process are still not fully understood^36–38^.

We aimed to generate T4 phage variants carrying catalytically inactive versions of Alt or ModA by exchanging single amino acids essential for their ADP-ribosyltransferase activity^39^. Based on the previous studies, we generated Alt E577A and ModA E165A mutants and confirmed them to be catalytically inactive *in vivo* (Fig. 3a, Extended Data Fig. 4).

**Fig. 3:**
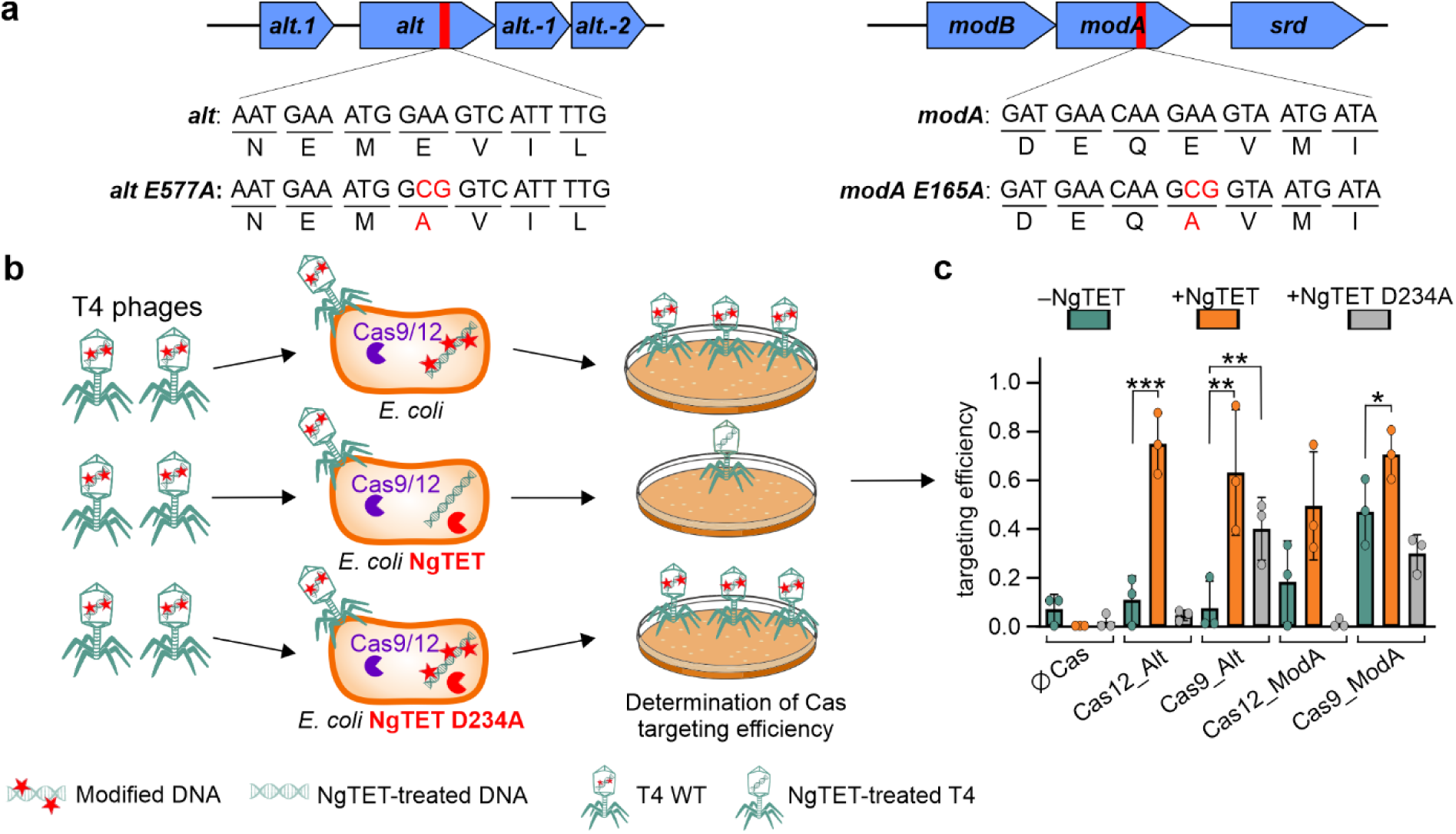
Effect of NgTET overexpression on CRISPR-Cas targeting efficiency *in vivo*. **a,** Target mutations in *alt* and *modA* genes that lead to the abolishment of ADP-ribosylation activity. **b,** Design of TE experiment. **c,** Evaluation of Cas-mediated T4 DNA TE *in vivo* in the presence or absence of NgTET/NgTET D234A (two-sided Student’s t-test, * - *P*_signif_ < 0.05, ** - *P*_signif_ < 0.025, *** - *P*_signif_ < 0.0125). *n =* 3 biological replicates.

To generate T4 phages carrying Alt E577A and ModA E165A mutations, we employed the heterologously expressed CRISPR-Cas9 and Cas12 systems, which were previously evaluated for the application to engineer phage T4^15,35^. For both CRISPR-Cas systems, the targeting of WT T4 DNA was reported to be strongly hindered by phage DNA modifications, leading to overall low mutagenesis efficiency^15,35^.

To introduce the site-specific point mutations into the phage DNA, we designed the spacers to precisely target the intended mutation insertion sites within *alt* and *modA*. We conducted targeting efficiency (TE) tests to evaluate Cas nucleases’ efficiency in targeting phage DNA. Therefore, we infected an *E. coli* strain that heterologously expressed one of the analyzed CRISPR-Cas systems with a spacer designed to target the *alt* or *modA* gene with a specific number of T4 phages. To determine the number of phages capable of efficiently infecting the host in the presence of CRISPR-Cas, we conducted a plaque assay (Fig. 3b). The number of plaques formed on the plate represented the phages that successfully evaded CRISPR-Cas targeting and propagated effectively. Based on this number, we calculated the reduction in phage numbers due to CRISPR-Cas targeting, which corresponds to the TE for the specific CRISPR-Cas system and the spacer.

First, we determined for each construct TE after T4 infection of wild-type *E. coli* expressing respective CRISPR-Cas systems (Fig. 3b). In this case, the 5ghmdC modifications were still present on >99% of T4 cytosines (Fig. 1d), potentially hindering the activity of CRISPR-Cas systems. In the absence of NgTET, most constructs showed a TE of 0 to 0.1, except for Cas9_ModA, which showed a TE of ∼0.5. These results confirm the generally low targeting efficiency of T4 WT DNA by CRISPR-Cas9 and Cas12, which is additionally influenced by the specific site targeted within the phage genome, as also reported in earlier studies^15,35^.

Next, we investigated whether the presence of NgTET enzyme, leading to a reduction of DNA modifications on T4 DNA, can increase the Cas targeting efficiency of T4 DNA *in vivo*. We infected *E. coli* recombinantly expressing the respective CRISPR-Cas system and NgTET simultaneously. The expression of NgTET increased the TE for all analyzed targets (Fig. 3c). For example, for CRISPR-Cas12 targeting *alt*, we observed a significant 7-fold increase in TE from 0.11 to 0.74. As for CRISPR-Cas9 targeting *modA*, which already had a high TE before NgTET treatment (TE of 0.47), the TE increased by 1.5-fold due to NgTET treatment of DNA. To confirm the connection between the increased Cas cleavage activity on T4 DNA and altered DNA modification levels associated with NgTET activity, we performed a parallel experiment using the inactive mutant NgTET D234A for TE determination^32^. As expected, the expression of the inactive NgTET mutant did not improve targeting efficiency in all analyzed settings (Fig. 3c).

In summary, the NgTET dioxygenase activity on T4 DNA directly enhances its TE by CRISPR-Cas9 and Cas12 and strongly diminishes the TE’s reliance on the specific target site within the phage genome. This affords flexibility in spacer design and the ability to target the desired genome region, which is a critical factor for mutagenesis and the introduction of SNPs into the modified genome.

### Temporarily removal of T4 DNA modifications enables efficient phage T4 mutagenesis

The treatment of T4 DNA with NgTET significantly enhances its targeting efficiency by CRISPR-Cas9 and -Cas12 systems (Fig. 3c). To validate that NgTET treatment indeed enhances phage mutagenesis through improved CRISPR-Cas targeting, we conducted mutagenesis experiments on both the *alt* and *modA* genes, using T4 WT or NgTET-treated T4. To prove the hypothesis, we employed four distinct mutagenesis setups: (1) using CRISPR-Cas9 alone, (2) the CRISPR-Cas9 combined with NgTET dioxygenase, (3) CRISPR-Cas12 alone, and (4) CRISPR-Cas12 combined with NgTET dioxygenase.

In the settings where NgTET dioxygenase was expressed during the mutagenesis (settings 2 and 4), the phages were pretreated with NgTET before infecting the mutagenesis strain (Fig. 4). This pretreatment aimed to reduce the abundance of 5ghmdC and enhance the targeting of phage DNA by CRISPR-Cas, as demonstrated for TE experiments previously (Fig. 3c). Nevertheless since the TEs for both CRISPR-Cas systems and genes exhibited variability ranging between 0.5 and 0.75 (Fig. 3c), we anticipated that a fraction of the phages used for mutagenesis would evade CRISPR-Cas targeting. Consequently, mutagenesis was expected to yield a mixed population of wild-type and mutant T4 phages.

**Fig. 4:**
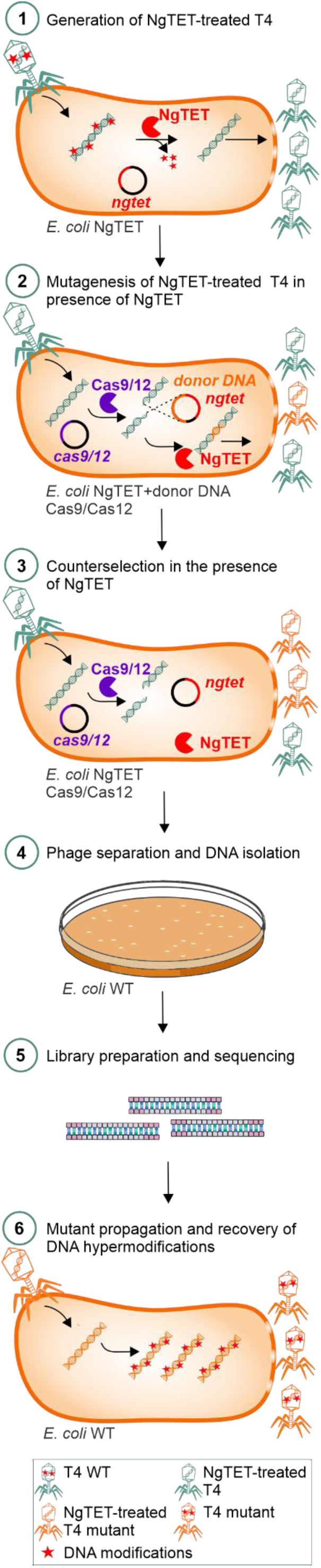
Established workflow for T4 phage mutagenesis and mutants screening.

As the NgTET treatment increases DNA susceptibility to specific CRISPR-Cas targeting, it suggests a potential strategy for enhanced counterselection with the same CRISPR-Cas system used for the mutagenesis. Accordingly, we conducted a counterselection round in the absence of donor DNA.

Following mutagenesis, the resulting phage population was isolated by plating with *E. coli* B strain (wild-type) and then screened for mutations, enabling the identification of specific genetic changes.

As mentioned above, a mixed population of the phages (wild-type and mutant phages) was expected. Although reporter genes could be introduced along the mutations to streamline mutant detection, they can potentially impact phage physiology, as outlined in the introduction. To avoid it, we adopted a multiplexed ONT-based approach^40^ for high-throughput screening of phage mutants. The mutation region was amplified from the phage with plaque-specific barcodes, which enables simultaneous screening of thousands of phages. Combining the mutagenesis with ONT screening, we determined that CRISPR-Cas9 and Cas12-based mutagenesis in the absence of NgTET did not result in the introduction of point mutations for all tested CRISPR-Cas systems, spacers, and targeted genes (Table 1). However, expression of NgTET in the infected *E. coli* host strain boosted the mutagenesis efficiency and resulted in 2.6-5.6% positive hits for targeted *modA* and *alt* genes (Extended Data Fig. 5-6). Here, the utilization of the CRISPR-Cas12 combined with NgTET resulted in successful mutagenesis of both analyzed genes, while the application of the CRISPR-Cas9 in combination with NgTET yielded the desired mutant only for *modA* gene.

**Table 1:** Summary of phage genome editing efficiency in the presence and absence of NgTET (in green: conditions resulting in positive clones, n=1)

| Edit name | Edited gene | Mutagenesis strain (*E. coli* BL21(DE3)...) | Plaques screened | Mutants identified | Mutation success rate (%) |
| --- | --- | --- | --- | --- | --- |
| *alt E577A* | *alt* | *alt* E577A, Cas9_Alt | 47 | 0 | 0 |
|  |  | **NgTET_***alt* E577A, Cas9_Alt | 45 | 0 | 0 |
|  |  | *alt* E577A, Cas12_Alt | 38 | 0 | 0 |
|  |  | **NgTET_***alt* E577A, Cas12_Alt | 44 | 1 | 2.6 |
| *modA E165A* | *modA* | *modA* E165A, Cas9_ModA | 36 | 0 | 0 |
|  |  | **NgTET_***modA* E165A, Cas9_ModA | 36 | 1 | 2.8 |
|  |  | *modA* E165A, Cas12_ModA | 35 | 0 | 0 |
|  |  | **NgTET_***modA* E165A, Cas12_ModA | 36 | 2 | 5.6 |

In conclusion, coexpressing NgTET dioxygenase in the mutagenesis strain to reduce the T4 DNA modification levels and enhance DNA targeting by Cas nucleases (Fig. 4), efficiently facilitated the introduction point mutations to phage T4 genome. These mutants could not be generated in the experimental setup without NgTET. Therefore, NgTET treatment of phage DNA eliminates the need for screening for the most efficient spacer, enabling the targeting of any desired position within the phage genome and the introduction of point mutations.

### Epigenetic cytosine modifications are common in phage genomes

To evaluate the applicability of the NgTET system for phage engineering beyond phage T4, we examined the occurrence of homologs of T4 DNA-modifying enzymes in other phages. We identified a total of 494 homologs for T4 gene 42, 131 for α-glycosyl-transferase, and 40 for β-glycosyl-transferase (Fig. 5, Supplementary Table 1).

**Fig. 5:**
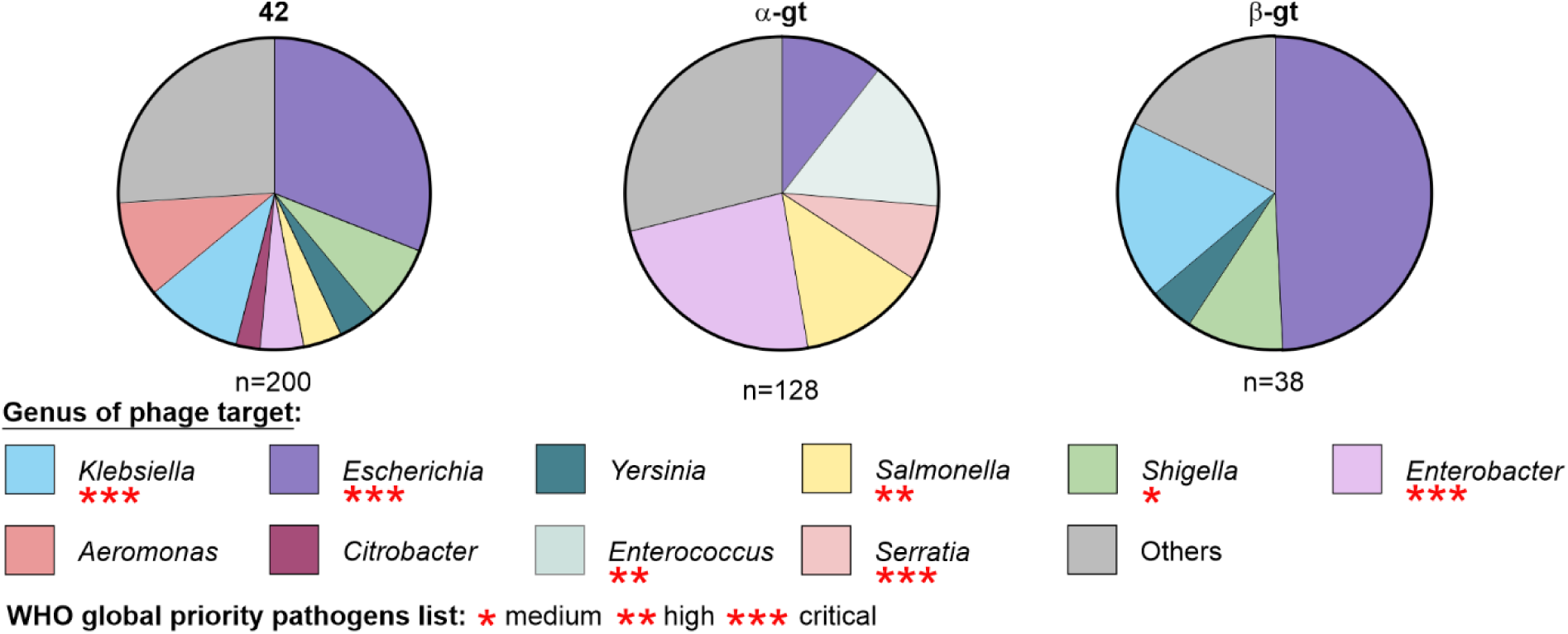
Distribution of cytosine modifying enzymes homologous to T4-originating enzymes 42, α-gt and β-gt among phages based on the genus of infected bacteria (BLAST score >80). Red asterisks highlight the global priority pathogens classification of the bacterial genus according to the WHO.

The prevalent identification of the homologous of gene 42 responsible for the initial hydroxymethylation of dC to 5hmdC suggests the presence of related modifications in several other phages, potentially involving glycosylations, arabinosylations, and other sugar modifications^18,41,42^. Notably, homologous enzymes were found in phages infecting a broad spectrum of bacterial genera. A considerable number of these bacterial hosts are pathogenic and belong to World Health Organization (WHO) priority strains^43^. Therefore, given the widespread presence of homologous DNA modifiers among phages and the potential of NgTET treatment demonstrated in this study on phage T4 DNA — both for investigating the role of DNA modifications and for efficient phage mutagenesis — the application of NgTET could be expanded to other phages. This extension would hold significant promise for both fundamental research and phage engineering.

## Discussion

CRISPR-Cas has emerged as an extremely valuable tool for genome engineering. The effectiveness of CRISPR-Cas systems in phage engineering has been limited by extensive DNA modifications up to this point. In particular, specific Cas-mediated targeting and induction of DNA double-strand breaks, essential for the mutagenesis, is prevented by DNA modifications^15,25^. The low Cas targeting efficiency of phage DNA is further reflected in the low mutagenesis rate, necessitating the screening of a large phage population to identify the phage mutants. Our study circumvents previous limitations and presents a novel approach for precise, site-specific phage mutagenesis based on temporarily reducing phage DNA modifications. This enhances DNA accessibility for Cas nucleases, boosting the efficiency of phage DNA double-strand breaks – the first and essential step of CRISPR-Cas-based mutagenesis. Thus, we eliminate the need for pre-screening for the most efficient spacer, significantly facilitating the introduction of scarless mutations at selected positions within the phage genome.

Moreover, the phage mutagenesis efficiency of up to 6% reported in this study, combined with the ONT-based high-throughput screening method, simplifies the detection of the point mutations within the phage genome, thereby reducing the need to screen large phage populations. This approach avoids additional genomic changes associated with reporter gene insertion and the potential drawbacks on phage DNA packaging^10^, allowing for the efficient, scarless introduction of point mutations into the phage genome.

To demonstrate the feasibility of our approach, we successfully introduced point mutations to inactivate T4 ARTs Alt and ModA. ARTs have been shown to be important for efficient host hijacking, which is achieved by introducing post-translation protein modifications such as ADP-ribosylation and RNAylation^44^. In addition to its catalytic function, Alt has been reported as an important T4 phage structural protein^7^. Therefore, deleting the entire gene would not solely allow the study of ADP-ribosylation’s role, but might also have a more pronounced impact on the phage infection. Thus, introducing a point mutation via our NgTET tool to inactivate the enzyme while preserving its role as a structural protein facilitates the functional characterization of the enzyme.

The modulation of T4 phage DNA modifications by NgTET treatment is possible due to the decoupled introduction of hydroxymethylation and glycosylation on T4 cytosines. The biosynthesis of modifications within phage T4 DNA occurs at both the nucleotide and polynucleotide chain levels. Initially, the hydroxymethylation is formed on dCMP, which is next converted to 5-hydroxymethylated 2’-deoxycytidine 5’-triphosphate and is integrated into phage DNA during replication. The glycosylation of so-formed 5hmdC occurs at the DNA level^18^. The fact, that glycosylation happens directly on the DNA, and not on the nucleotide levels, allows NgTET, which also acts on DNA level, to oxidize 5hmdC into 5fdC and 5cadC, thereby preventing the glycosylation. The propagation of the resulting phage progeny in the strain without NgTET, allows the glycosyltransferases to introduce the glycosylations back and thereby recover the wild-type-like modifications of phage T4 DNA. Such a recovery of DNA modifications in the subsequent phage generation highlights the major advantage of NgTET-coupled CRISPR-Cas mutagenesis over performing it with *α/β-gt* deletion or amber strains. In such deletion or amber strains, DNA glycosylations are permanently absent, making the phage consistently more susceptible to nucleases and thereby reducing its fitness. Additionally, NgTET treatment results in a fraction of non-modified 2′-deoxycytidines (approximately 35% for dC). To our knowledge, this was not possible by amber mutations or deletion of gene 42 due to its essentiality for phage infection^7,22^. Therefore, the presence of non-modified dC allows for efficient phage mutagenesis and also holds promise for studying the biological roles of various phage modifications or improving phage genome sequencing. Specifically - for sequencing approaches - NgTET treatment could be applied to reduce modification-mediated errors^45^, improving sequencing accuracy by decreasing the DNA modifications.

An important feature of the NgTET/CRISPR-Cas-coupled gene editing technique - presented in this study - is its potential transferability to other bacteriophages. Our homology search results show the cytosine-modifying enzymes being distributed among various phages (Fig. 5). Particularly, homologs of gene 42, responsible for the conversion of dC to 5hmdC, were detected in numerous phages. Gene 42 introduces a chemically active hydroxymethyl group to 2′-deoxycytidines, allowing further functionalization with various sugars and other functional groups. As a result, the modified DNA becomes impervious to degradation by DNA-binding nucleases, enabling the phages to effectively evade the host defense mechanisms. The capability of NgTET to act on the precursors 5mdC and 5hmdC, generated by homologous of gene 42, broadens the utility of our approach to engineer and to study phages carrying such DNA modifications. Notably, such phages are known to infect clinically and biotechnologically relevant bacterial genera, including *Klebsiella*, *Salmonella*, and *Serratia*. Therefore, NgTET-mediated mutagenesis has the potential to become a valuable tool for studying such phages by facilitating their mutagenesis, and taking us a step closer to the generation of "designer phages".

Lastly, it is worth considering the possibility of homologous enzymes to eukaryotic TET dioxygenases existing within bacteria. These enzymes may potentially be involved in counteracting phage infections through mechanisms analogous to those synthetically applied in this study. Reducing the extent of DNA modifications could enable nucleolytic cleavage of the invader’s DNA, which would be a powerful anti-phage defense system. On the other hand, in the case of phages, homologs of TET dioxygenases have been observed^46^. Yet, their involvement is typically associated with the modification of 2′-deoxycytidines, involving oxidation from 5mdC to 5hmdC. In contrast to NgTET, no following oxidation to 5fdC, 5cadC and dC has been described to date.

Taken together, the field of bacteriophage epigenetics remains widely unexplored and, therefore, represents an intriguing subject for future research, providing the opportunity to gain profound insights into the ongoing arms race between bacteria and bacteriophages.

## Methods

### Cloning of NgTET, sgRNAs and sequences for homologous recombination

The gene encoding for the NgTET from *Naegleria gruberi* was purchased from IDT as a gblock and amplified by PCR. XhoI and NcoI restriction sites were introduced during the PCR amplification of the vector. The resulting PCR product was digested with XhoI and NcoI and introduced into the pET-28a vector (Merck Millipore). The ModA E165A, Alt E577A, and NgTET D243A were generated by site-directed mutagenesis. The insertion of the sgRNA sequences into pCPf1 (#122185, Addgene) and DS-SPCas (#48645, Addgene) plasmids was performed via complete plasmid amplification with the primers carrying the respective 5’-overhangs. The linearised plasmid was circularised via blunt-end ligation. All resulting plasmids were sequenced by Sanger sequencing and transformed into chemically competent *E. coli* BL21 (DE3). All primers and strains used in this work can be found in Supplementary Table 2. The plasmid maps are deposited at https://github.com/MaikTungsten/CRISPRT4

### Phage T4 propagation

For phage T4 propagation, *E. coli* BL21 (DE3) cells were used. The initial culture was set to OD_600_ ∼0.1 and grown at 37°C, 160 rpm until OD_600_ ∼0.8 was reached. Phage T4 was added to a multiplicity of infection (MOI) of 0.8 together with 1 mM MgCl_2_ and 1 mM CaCl_2,_ and the infection was run for 3 – 4 h at room temperature, 120 rpm. The lysate was centrifuged at 1,200 x *g* for 5 min and the supernatant was filtered through Steritop filters (pore size 0.45 µm). The concentration of the phages was determined via plaque assay and the phage suspension was stored at 4°C.

For the propagation of NgTET-treated T4 phages, the infection was performed in the same way using *E. coli* BL21 (DE3) pET28a_NgTET. The medium was supplied with 50 µM kanamycin. As the culture reached OD_600_ ∼0.4, expression of NgTET was induced by the addition of 50 µM Isopropyl-β-d-thiogalactopyranosid (IPTG). The culture was grown for another 2 h at 37°C and 160 rpm and infected with phage T4 as described above. To recover NgTET-treated T4, the infection of *E. coli* BL21 (DE3) was repeated with NgTET-treated T4 as described above.

### Plaque assay and efficiency of plaquing

*E. coli* culture of interest was grown to OD_600_ ∼0.8 in the presence of strain-specific antibiotics (Supplementary Table 2). Subsequently, 300 µL of the culture were infected with T4 WT phage or NgTET–treated T4 phage with either defined or unknown MOI. The bacteria-phage suspension was incubated at 37°C for 7 min, followed by transfer to 4 ml of LB-soft-agar (0.75%), thoroughly mixed. After mixing, the suspension was poured onto an LB-agar plate. According to the respective strains, the agar was supplemented with corresponding antibiotics. The plates were incubated at 37°C overnight and subsequently assessed for plaque-forming units (PFU) determination. To determine the efficiency of plaquing, the number of resulting plaque-forming units (pfu) in analyzed settings was divided by the input PFU.

### Growth and Lysis assays

The growth and lysis of the bacterial cells were assessed by measuring the optical density of the bacterial cultures at 600 nm. This was done either by manually withdrawing samples for OD_600_ measurements or using the Tecan Spark plate reader (Tecan Group, Männedorf, Switzerland). To determine the growth curves, initial cultures were inoculated at an OD_600_ of 0.1, and the growth was monitored until the stationary growth phase was reached. For the lysis assays, the bacterial culture of interest was grown starting from OD_600_ 0.1, with the addition of antibiotics if necessary. In the case of NgTET expression, induction occurred at OD_600_ of 0.4 by adding 100 µM IPTG. Phage infection was carried out at OD_600_ of 0.8 with MOI of 3. The lysis was conducted at 23°C.

### Detection of inactive mutants of Alt and ModA

The *in vivo* activity of the ARTs mutants was assessed by expressing the target proteins in *E. coli* BL21 (DE3) strains of interest (Supplementary Table 2). To confirm the presence of ADP-ribosylation events, the cells were grown at 37°C and were sampled first at OD_600_ of 0.8 for Alt and 1.2 for ModA. Afterwards, the induction was performed with 1 mM IPTG, and proteins were expressed at 37°C (for Alt and its mutants) and 4°C (for ModA and its mutants). After 1 h of expression, the cultures were sampled again and submitted to 12% SDS-PAGE. For protein visualization prior to blotting, the gels were supplemented with 5%(v/v) 2,2,2-trichloroethanol (TCE) and proteins were visualized under UV-transillumination (300 nm) for 60 s^47^. The gels were then equilibrated in transfer buffer (25 mM Tris (pH 8.3), 192 mM glycine, 0.1% (v/v) SDS, and 20% (v/v) methanol), and proteins transferred to nitrocellulose membrane (NCM) in a semi-dry manner at 400 mA for 50 min. After blotting, membranes were washed 3 times with TBS-Tween (TBS-T; 10 mM Tris-HCl, pH 7.5, 150 mM NaCl, 0.05% [v/v] Tween® 20). Subsequently, the membranes were blocked in 5% [w/v] milk powder in TBS-T for 1 h at room temperature. For the detection of ADP-ribosylated proteins, the membranes were incubated o/n at 4°C in 10 ml washing buffer (1% [w/v] milk powder in TBS-T) containing a 1:10,000 dilution of anti-pan-ADP-ribose binding reagent MABE1016 (Merck) at 4°C^48^. After washing, the membranes were incubated with 10 ml of a 1:10,000 dilution of horseradish-peroxidase-(HRP)-goat-anti-rabbit-IgG secondary antibody (Advansta) in washing buffer at room temperature for 1 h. Following another wash with PBS, the ADP-ribosylated proteins were visualized using chemiluminescence using the SignalFire ECL Reagent or the SignalFire Elite ECL Reagent (Cell Signaling Technology) according to the manufacturer’s instructions.

### Phage DNA isolation

Phage suspension of interest with a concentration of >10^10^ PFU/ml was pretreated with 20 U DNase I (ThermoFisher Scientific, MA, USA) and 2 µL RNase A/T1 Mix (4 µg RNase A, 10 U RNase T1, ThermoFisher Scientific, MA, USA) at 37°C for 30 min to remove host-originating nucleic acids. Next, the phage was purified in a 0-45% sucrose gradient, generated in TM buffer (50 mM Tris-HCl, 10 mM MgCl_2_, pH 7.5). Therefore, 500 µL of the phage solution was loaded on top of the gradient and centrifugated at 70,000 x *g*, 20 min, 4°C. The fraction of the gradient containing the phages was removed with a blunt cannula and transferred into a new ultracentrifugation tube. 30 ml of ice-cold TM buffer were added and the phages were pelleted at 100,000 x *g* for 1 h at 4°C. The supernatant was discarded and the pellet was resuspended in 500 µL TM buffer and incubated at 4°C overnight. 1 µg of Proteinase K (Carl Roth, Karlsruhe, Germany) was added and the samples were incubated for another 30 minutes at 37°C. For DNA isolation, a phenol/chloroform/isoamyl alcohol (P/C/I) (Carl Roth, Karlsruhe, Germany) extraction was performed three times. The remaining phenol was removed by chloroform back-extraction (3x). The DNA was precipitated with 0.1 volume of 3 M NaOAc (pH 5.5) and 2.5 volumes of absolute ethanol at −20°C overnight. The DNA was pelleted at 15,000 x *g* at 4°C for 1 h and the pellet was washed twice with 70% ethanol. The purified DNA was resuspended in Millipore water and stored at −20°C until further use. For LC-MS analysis, the isolated DNA was processed to single nucleosides by applying Nucleoside Digestion Mix (NEB, MA, USA).

### Phage T4 mutagenesis

*E. coli* pET-28a_NgTET_*x* + pCpf1_*x*/DS_SPcas_*x* (x: donor DNA and respective sgRNA) strains were used for mutagenesis (Supplementary Table 2). The cells were cultivated at 37°C and 160 rpm until OD_600_ ∼0.4, after which NgTET expression was induced by the addition of 50 µM IPTG. The cultivation was proceeded until OD_600_ of 0.8. At this point, the cultures were adjusted to room temperature and 120 rpm and supplemented with 1 mM MgCl_2_ and 1 mM CaCl. The cells were infected with T4 NgTET phage set to MOI of 0.5. The infection was performed for 3 h, 130 rpm, and 23°C after which the cells were pelleted and supernatant filtered through Steritop filters (pore size 0.22 µm). The titer of the newly generated phages was determined via plaque assay and the phages were used for the counterselection via infection of *E. coli E. coli* pET-28a_NgTET_*x* + pCpf1_*x*/DS_SPcas_*x* strain. The counterselection was performed under the same conditions as the mutagenesis. The counterselected phages were filtered and used for a plaque assay with *E. coli* B strain. Single plaques were picked and transferred to 100 µl Pi-Mg buffer (26 mM Na_2_HPO_4_, 68 mM NaCl, 22 mM KH_2_PO_4_, 1 mM MgSO_4_, pH 7.5) supplemented with 2% chloroform. The suspension was incubated for 1 h at room temperature and was further used for mutagenesis analysis via sequencing, infection of *E. coli*, or stored at 4°C until further use.

### LC-MS

Relative determination of dA, dT, dC, 5hmdC, 5fdC, 5cadC, and 5ghmdC was performed using HRES LC-MS. The chromatographic separation was performed on a ThermoFisher Scientific Vanquish HPLC System using a Atlantis T3 C18 column (150 x 2.1mm, 100 A, 3 µm, Waters, MA, USA) equipped with a 20 × 2.1 mm guard column of similar specificity at a constant eluent flow rate of 0.2 ml/min and a column temperature of 40 °C with eluent A being 10 mM Ammonium Acetate in water at a pH of 4.5 and eluent B being 0.1 % of formic acid in MeOH (Honeywell, NC, USA). The injection volume was 5 µl for standards and T4 WT samples and 20 µl for NgTET-treated samples.

The mobile phase profile consisted of the following steps and linear gradients: 0 – 1 min constant at 5% B; 1 – 5 min from 5 to 90% B; 5 – 7 min constant at 90% B; 7 – 7.1 min from 90 to 5% B; 7.1 to 12 min constant at 5% B.

A Thermo Scientific ID-X Orbitrap mass spectrometer was used in negative and positive ionization mode (separate injections) with a High-temperature electrospray ionization source and the following conditions: H-ESI spray voltage at 3400 V(+), 2400 V (-) sheath gas at 35 arbitrary units, auxiliary gas at 7 arbitrary units, sweep gas at 0 arbitrary units, ion transfer tube temperature at 300°C, svaporiser Temperature at 275°C.

Detection was performed in full scan mode using the orbitrap mass analyzer at a mass resolution of 120 000 in the mass range 200 – 450 (m/z).

Extracted ion chromatograms of the [M-H]^-^ (dA, dT, dC, 5hmdC, 5fdC, 5cadC)/[M+H]^+^ (5ghmdC) forms were integrated using Tracefinder software (ThermoFisher Scientific, MA, USA). Relative abundance in each sample was calculated by normalizing the peak area of each peak by the peak area of the dG signal in each specific sample, thus using dG as a sample-specific internal standard.

### Screening for phage T4 mutants by multiplexed Nanopore amplicon sequencing

The method for highly multiplexed Nanopore sequencing is based on Ramírez Rojas *et al*. 2024 ^40^. Briefly, the method relies on a series of two PCRs to generate the multiplex amplicon DNA for Nanopore sequencing. The first PCR attaches standardized overhangs (M13 fwd/rev sequences in this instance), serving as an amplification sequence for the second PCR attaching the barcodes. 1 µL of isolated phage T4 in Pi-Mg buffer was used as a PCR template in the first PCR with the following reaction mix: 0.125 µM each primer (Supplementary Table 2) in 1x High Fidelity Master Mix (NEB) with a total volume of 10 µl. PCR settings: 98 °C for 30 s followed by 30 cycles 98 °C 20 s, 69 °C 30 s, and 72 °C 2 min with a final hold at 72 °C 5 min. The dual barcodes were attached in a second PCR with the following reaction mix using KAPA HiFi HotStart ReadyMix (Roche), using 1 µL of a 1:10 dilution of the initial PCR as a template and 0.3 µM of the barcoding primers (Supplementary Table 2) in 7 µl reactions. PCR settings: 95 °C for 3 min followed by 20 cycles 98 °C 20 s, 66 °C 15 s, and 72 °C 60 s with a final extension 72 °C 5 min at hold at 12 °C. All the barcoded PCR reactions were pooled and purified using NucleoMag kit (Macherey Nagel) for NGS library preps. Briefly, DNA was bound to magnetic beads, washed twice with 80% ethanol, and eluted in 100 µL elution buffer (5 mM Tris-HCl pH 8.5). Concentration was determined with Nanodrop (ThermoFisher Scientific) and Qubit (Invitrogen) using the broad range and/or high-sensitivity assay. Sequencing libraries were generated with the SQK-LSK109 Ligation Sequencing kit (Oxford Nanopore Technologies) according to the manufacturer guidelines starting with 1 µg of input DNA. Sequencing was performed on Flongle flow cells (R9.4.1 chemistry) on a MinION device.

### Analysis of long-read sequencing data

Nanopore raw reads were basecalled using guppy (v6.1.2 to v6.4.2), basecalled raw reads are deposited under BioProject PRJNA952186. In a miniconda environment, reads were demultiplexed using minibar^49^ and mapped to the phage T4 reference genome (NC_000866.4) using minimap2 (version 2.24)^50^. The resulting SAM files were converted to BAM files, sorted, and indexed with samtools (version 1.4.1)^51^. Variant calling in the target region for mutagenesis was subsequently performed using longshot (version 0.4.1)^52^ and resulting VCF files were inspected for desired point mutants with Integrative Genomics Viewer (IGV 2.16.0)^53^. Further, read counts per sample and target region were obtained from sorted BAM files using featureCounts (subread, version 2.0.1)^54^ and a custom R script. Detailed code and an easy command line application for these data analysis steps are provided to the community with CRISPRT4 (code available: https://github.com/MaikTungsten/CRISPRT4)

### Phylogenetic analysis

The identification of homologs for proteins 42, α-, and β-glycosyltransferases was conducted through a protein BLAST search against the NCBI database (data collected in May 2023) (Supplementary Table 1). Only sequences originating from phages were selected for further analysis to maintain specificity.

## Acknowledgments

We thank Tristan Krause, Petra Mann, Ruzica Sedic and Madita Viering for their experimental assistance. K.H. is supported by funding from the Max Planck Society and the German Research Council (DFG SPP 2330, Project number 464500427). A.A.R.R. and D.S. are supported by the Max Planck Society within the framework of the MaxGENESYS project. M.W.S. is supported by funding from the Joachim Herz Foundation and the Studienstiftung des deutschen Volkes e.V.

## Author contributions

K.H. and Na.P. designed the study and wrote the initial draft. F.A.B, Na.P. cloned, expressed, analyzed the NgTET system, and prepared samples for LC-MS. Na.P. established a mutagenesis pipeline, evaluated experimental results. Ni.P. and Na.P. established an LC-MS pipeline to analyze phage DNA composition. A.A.R.R. and D.S. performed library preparation and ONT-sequencing. M.W.S. established the pipeline for the evaluation of the sequencing results. All authors reviewed and edited the manuscript.

## Data availability statements

https://github.com/MaikTungsten/CRISPRT4

## Code availability statements

https://github.com/MaikTungsten/CRISPRT4

