## Supplementary Information for "Temporal epigenome modulation enables efficient bacteriophage engineering and functional analysis of phage DNA modifications"

Extended Data Fig. 1: Growth and lysis of *E. coli* upon different conditions.
Extended Data Fig. 2: LC-MS analysis of T4 DNA composition.
Extended Data Fig. 3: Effect of Cas expression on growth and lysis of *E. coli*.
Extended Data Fig. 4: Validation of Alt and ModA ARTs inactivation via Alt E577A and ModA E165A mutations.
Extended Data Fig. 5: Alt mutants sequencing summary.
Extended Data Fig. 6: ModA mutants sequencing summary.

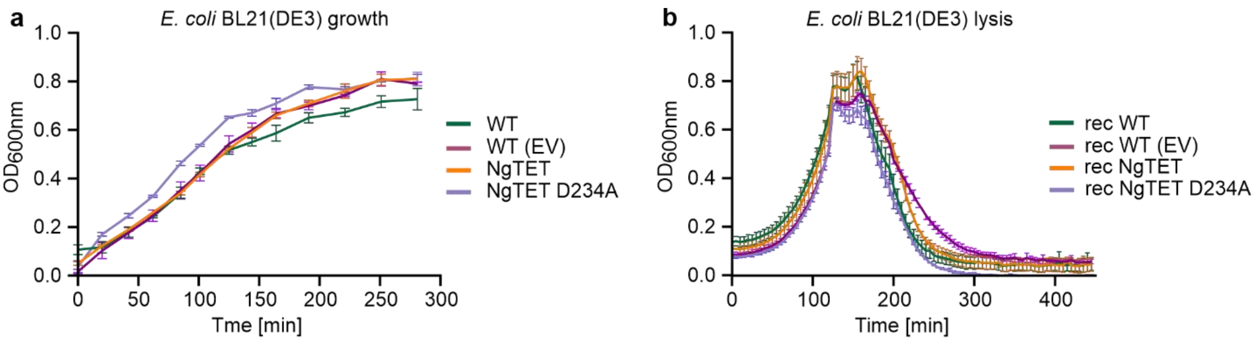


**Extended Data Fig. 1: Growth and lysis of *E. coli* upon different conditions. a,** Impact of NgTET recombinant expression on *E. coli* growth. *n* = 3 biological replicates. **b**, Lysis of *E. coli* by the phages recovered from different conditions. *n* = 3 biological replicates.

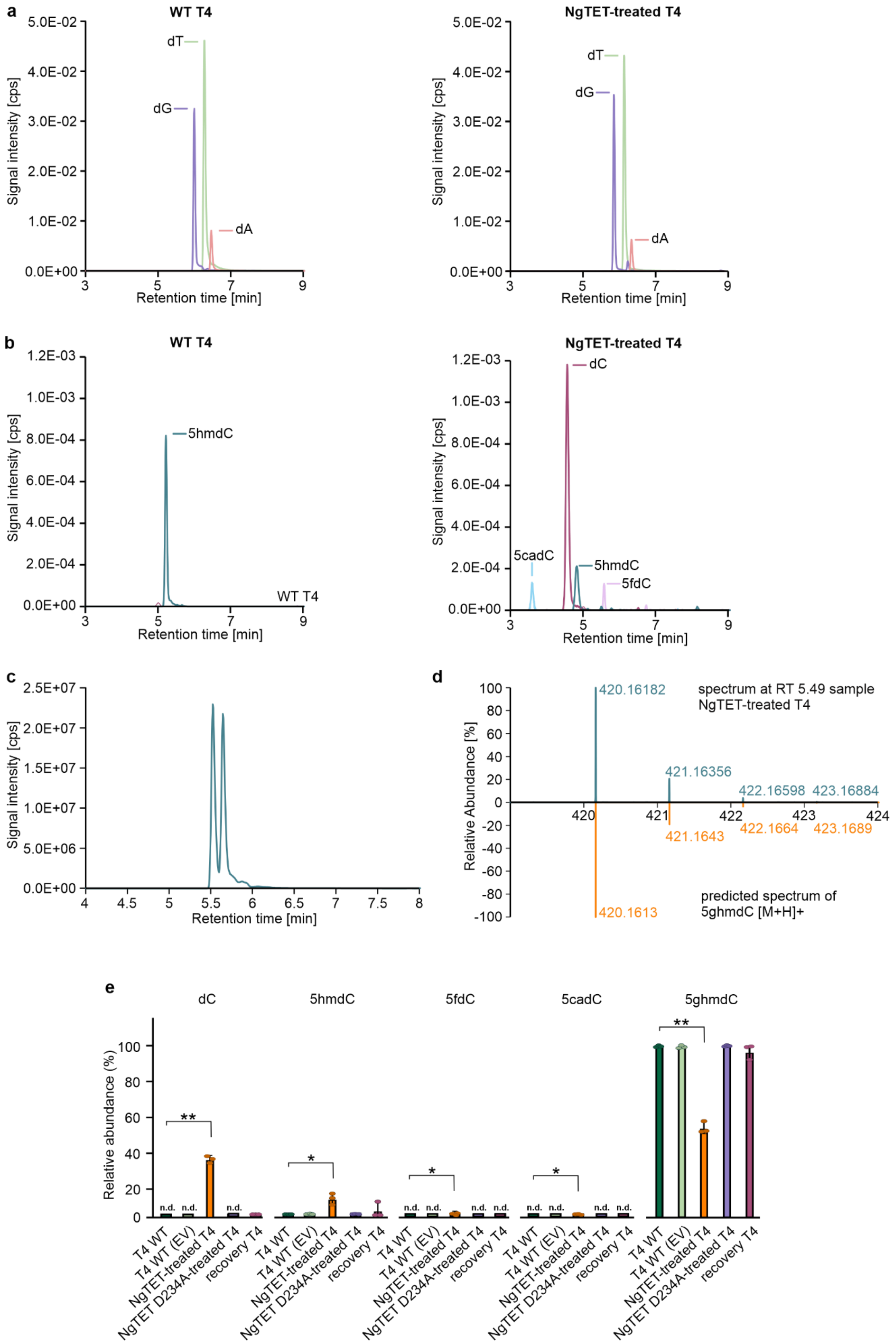
a
WT T4
NgTET-treated T4
Signal intensity [cps]
Retention time [min]
dT
dG
dA
b
WT T4
NgTET-treated T4
5hmdC
dC
5cadC
5fdC
c
d
spectrum at RT 5.49 sample NgTET-treated T4
420.16182
421.16356
422.16598
423.16884
421.1643
422.1664
423.1689
420.1613
predicted spectrum of 5ghmdC [M+H]+
Relative Abundance [%]
e
dC
5hmdC
5fdC
5cadC
5ghmdC
Relative abundance (%)
n.d.
T4 WT
T4 WT (EV)
NgTET-treated T4
NgTET D234A-treated T4
recovery T4

**Extended Data Fig. 2: LC-MS analysis of T4 DNA composition. a-b,** Extracted ion chromatogram overlay of nucleotide mass traces in T4 WT DNA (front) and NgTET-treated T4 DNA (back). Data exemplifies differences in relative abundances of cysteine nucleotides (dC, 5hmdC, 5fdC, 5cadC) (b) while dG, dT and dA (a) show similar trends. Signal intensities have been normalized against the overall nucleotide signal in the respective sample to correct for differences in sample concentration and injection volume. **c-d,** Extracted ion chromatogram of presumed 5ghmdC (c). The fragmentation pattern of presumed 5ghmdC corresponds to the predicted one (d). **e**, Relative abundance of cytosine derivatives in T4 phage isolated from different strains (two-sided Student´s t-test, * - $P_{signif} < 0.05$, ** - $P_{signif} < 0.025$; n.d.: not detected). $n = 3$ biological replicates.

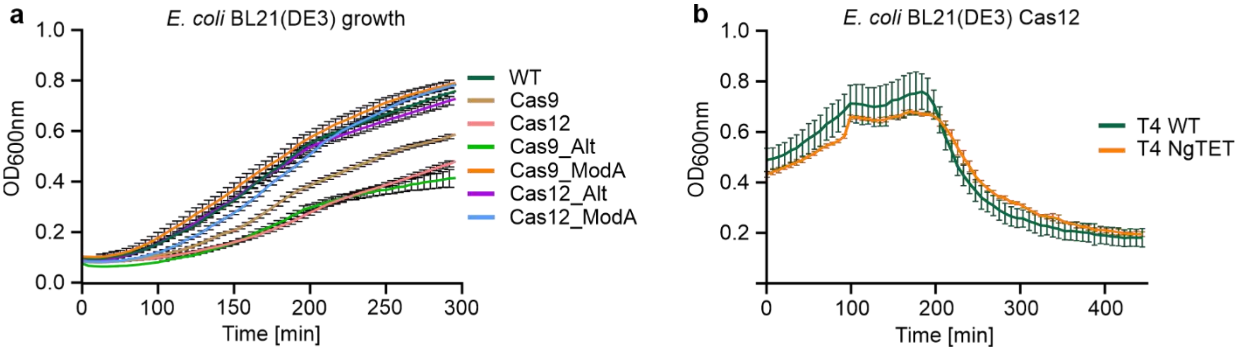


**Extended Data Fig. 3: Effect of Cas expression on growth and lysis of *E. coli*. a**, Impact of CRISPR/Cas9 or Cas12 systems recombinant expression on *E. coli* growth. *n* = 3 biological replicates. **b**, Impact of CRISPR/Cas12 system recombinant expression on T4 WT and T4 NgTET lysis efficiency. *n* = 3 biological replicates.

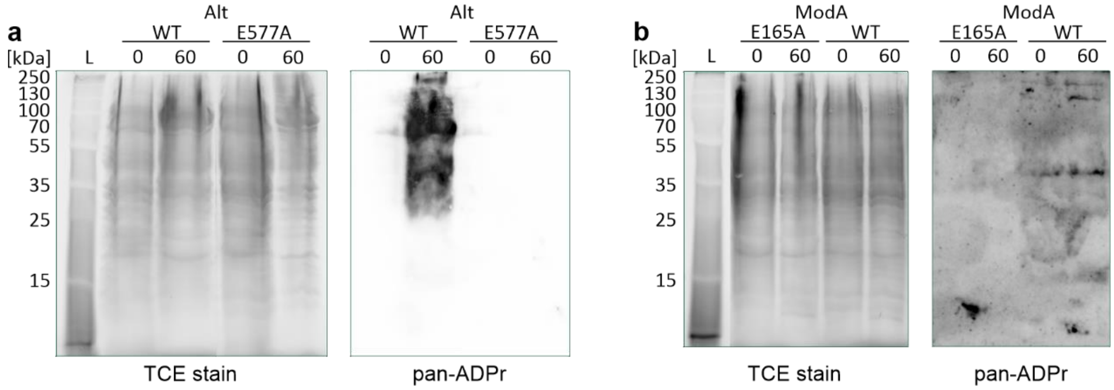


**Extended Data Fig. 4: Validation of Alt and ModA ARTs inactivation via Alt E577A and ModA E165A mutations. a-b**, Stain-free scan (TCE stain, loading control, left) and Western blot analysis (pan-ADPr antibody for ADP-ribosylation detection, right) to identify ADP-ribosylation events by ARTs and their mutants. Both mutants demonstrate complete abolishment of ADP-ribosylation. *n* = 3 biological replicates, a representative example is shown.

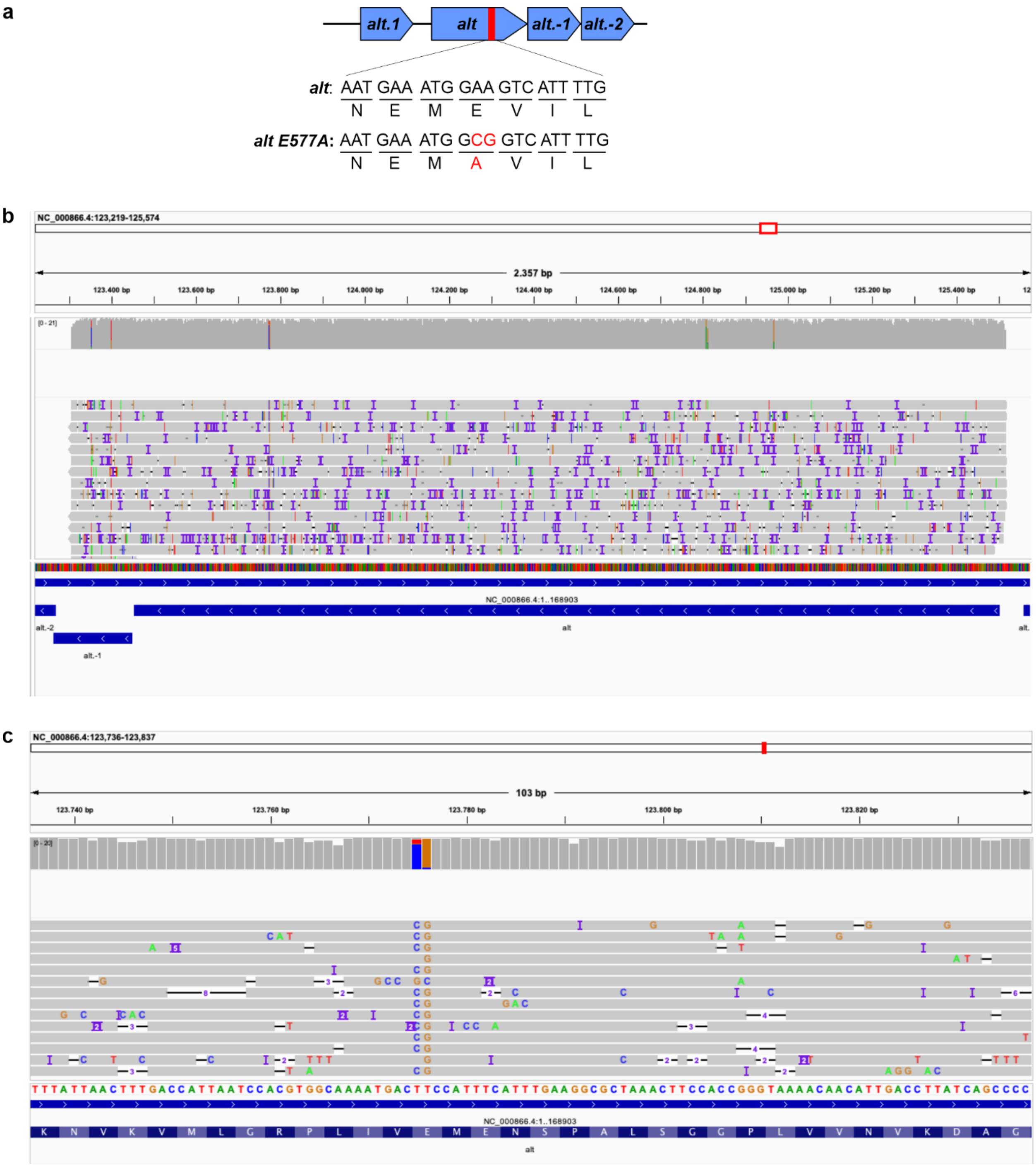


**Extended Data Fig. 5:** ***Alt*** **mutants sequencing summary. a**, The targeted mutation site in *alt* gene. **b**, Sequence coverage at the mutation site. **c**: Alt E577A mutant.

a

b

c

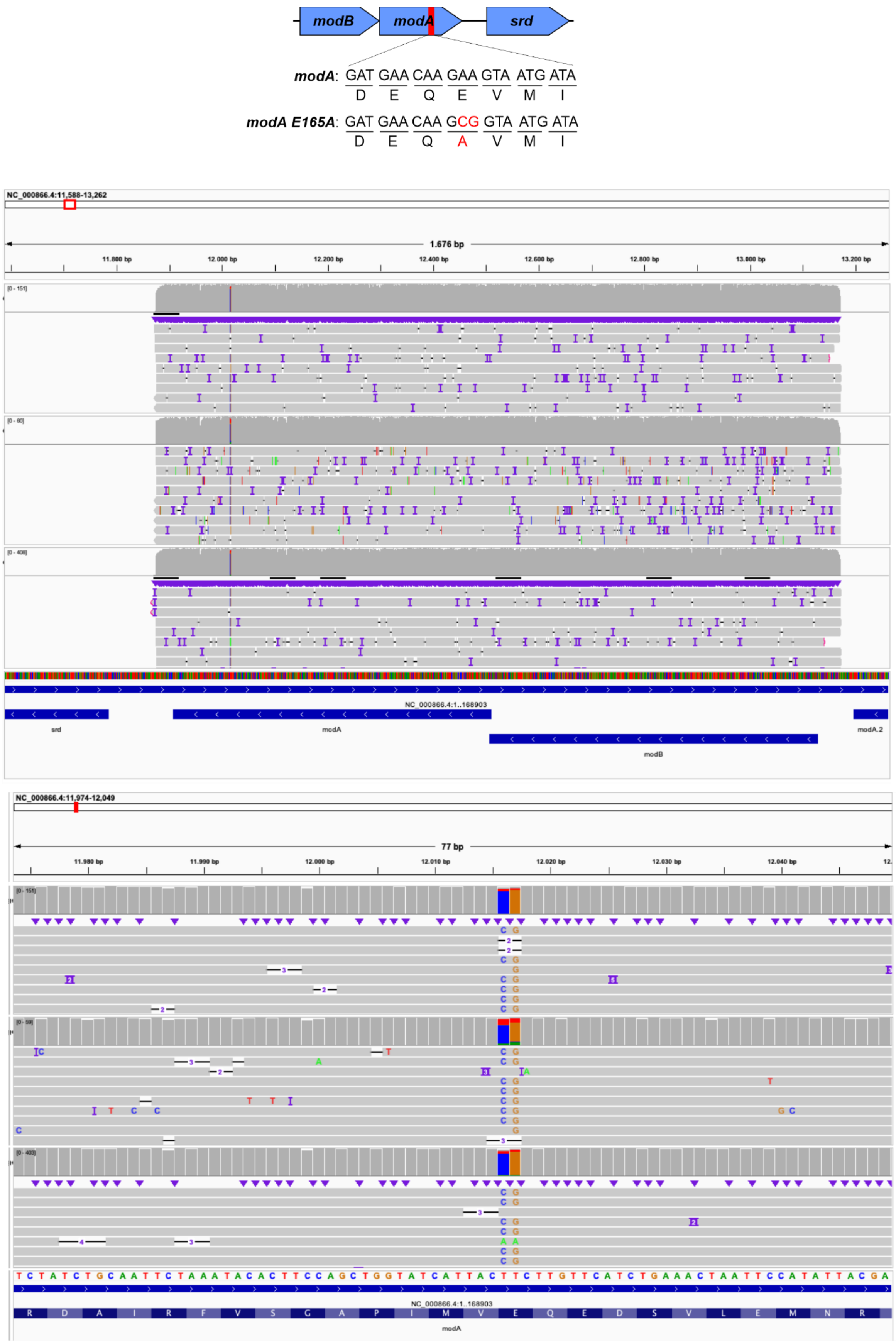

modB
modA
srd
modA: GAT GAA CAA GAA GTA ATG ATA
D E Q E V M I
modA E165A: GAT GAA CAA GCG GTA ATG ATA
D E Q A V M I
NC_000866.4:11,588-13,262
1,676 bp
11,800 bp
12,000 bp
12,200 bp
12,400 bp
12,600 bp
12,800 bp
13,000 bp
13,200 bp
NC_000866.4:1..168903
srd
modA
modB
modA.2
NC_000866.4:11,974-12,049
77 bp
11,980 bp
11,990 bp
12,000 bp
12,010 bp
12,020 bp
12,030 bp
12,040 bp
T C T A T C T G C A A T T C T A A A T A C A C T T C C A G C T G G T A T C A T T A C T T C T T G T T C A T C T G A A A C T A A T T C C A T A T T A C G A
NC_000866.4:1..168903
R D A I R F V S G A P I M V E Q E D S V L E M N R
modA

**Extended Data Fig. 6: *ModA* mutants sequencing summary. a**, The targeted mutation site in *modA* gene. **b**, Sequence coverage at the mutation site. **c**, ModA E165A mutant.
